# mInDel: an efficient pipeline for high-throughput InDel marker discovery

**DOI:** 10.1101/009290

**Authors:** Yuanda Lv, Yuhe Liu, Xiaolin Zhang, Han Zhao

## Abstract

**Background:** Next-Generation Sequencing (NGS) technologies have emerged as a powerful tool to reveal nucleotide polymorphisms in a high-throughput and cost-effective manner. However, it remains a daunting task to proficiently analyze the enormous volume of data generated from NGS and to identify length polymorphisms for molecular marker discovery. The development of insertion-deletion polymorphism (InDel) markers is in particular computationally intensive, calling for integrated high performance methods to identify InDels with high sensitivity and specificity, which would directly benefit areas from genomic studies to molecular breeding.

**Results:** We present here a NGS-based tool for InDel marker discovery (mInDel), a high-performance computing pipeline for the development of InDel markers between any two genotypes. The mInDel pipeline proficiently develops InDel markers by comparing shared region size using sliding alignments between assembled contigs or reference genomes. mInDel has successfully designed thousands of InDel markers from maize NGS data locally and genome-wide. The program needs less than 2 hours to run when using 20 threads on a high-performance computing server to implement 40G data.

**Conclusions:** mInDel is an efficient, integrated pipeline for a high-throughput design of InDel markers between genotypes. It will be particularly applicable to the crop species which require a sufficient amount of DNA markers for molecular breeding selection. mInDel is freely available for downloading at www.github.com/lyd0527/mInDel website.

## Background

For most plant species, genetic mapping of a trait of interest relies largely upon the trait-marker association revealed from segregating populations [1–3]. The quantity and quality of molecular markers are therefore instrumental to the resolution and accuracy of successful linkage construction, gene mapping, and cloning studies [4–7]. The development of a proficient and cost-effective marker system is also critical in other studies including population genetics, molecular evolution and marker assisted breeding, etc [8–10].

In the past decades, the Southern blot-based marker systems such as restriction fragment length polymorphism (RFLP) have been progressively replaced by PCR-based markers such as random amplified polymorphic DNA (RAPD), simple sequence repeat (SSR), and amplified fragment length polymorphism (AFLP). However, the development and large-scale screening of polymorphic molecular makers at the time usually involved extensive library construction and Sanger sequencing which are time-consuming and labor-intensive. The recent advent of next-generation sequencing (NGS) technologies has revolutionized the pace and throughput of sequence information generation, and greatly accelerated the discovery processes for genetic markers [11,12]. A large amount of single nucleotide polymorphisms (SNP) were developed based on NGS data and showed a strong vitality in several studies [11,13].

Compared to other PCR-based marker types and the more recent SNP system, Insertion-Deletion markers (InDels) have shown several advantages. Firstly, InDels are much more abundant than other PCR-based markers such as SSRs in genomes [15–18]. Secondly, InDels are potentially multi-allelic and codominant, offering more genomic information than the bi-allelic SNPs [18–20]. Lastly, existing InDels can be readily used to genotype new materials via simple gel electrophoresis, whereas SNP detection requires specialized and usually more expensive equipment (e.g. DHPLC) or assays (e.g. TaqMan). All these characteristics make InDel markers highly suitable for applications from basic genomic research to applied plant molecular breeding.

Faced with the great wealth of sequence information generated from NGS however, existing programs or pipelines for Indel marker discovery are struggling to keep in step. They are usually reliant on a reference genome and can only discover small indels (<= 10 nt) due to the limited alignment algorithm; the primer design efficiency is often compromised by the large nucleotide variation flanking the InDels, all of which inhibited their usage in plant molecular breeding. To overcome the aforementioned problems, here we present and validate an efficient and high-throughput pipeline for InDel markers development based on NGS data independent of a reference genome. The mInDel pipeline has been primarily designed to work with diploid plant genomes, but can also be applied to polyploid plant species such as cotton, wheat and potato.

## Implementation

### Overview of the pipeline

mInDel is a comprehensive and high performance marker development program to identify specific InDel primers between genotypes in a high-throughput manner. mInDel mainly implements five modules: the Pre-processing module, the *De novo* assembly module, the overlap PCR primer design in batch module, the ePCR mapping module and InDel screening and marker development module. In the following, we describe the mInDel pipeline for InDel marker development, including one pre-processing step, three main marker discovery steps and a post-processing step, as shown in Figure 1.

**Figure 1:**
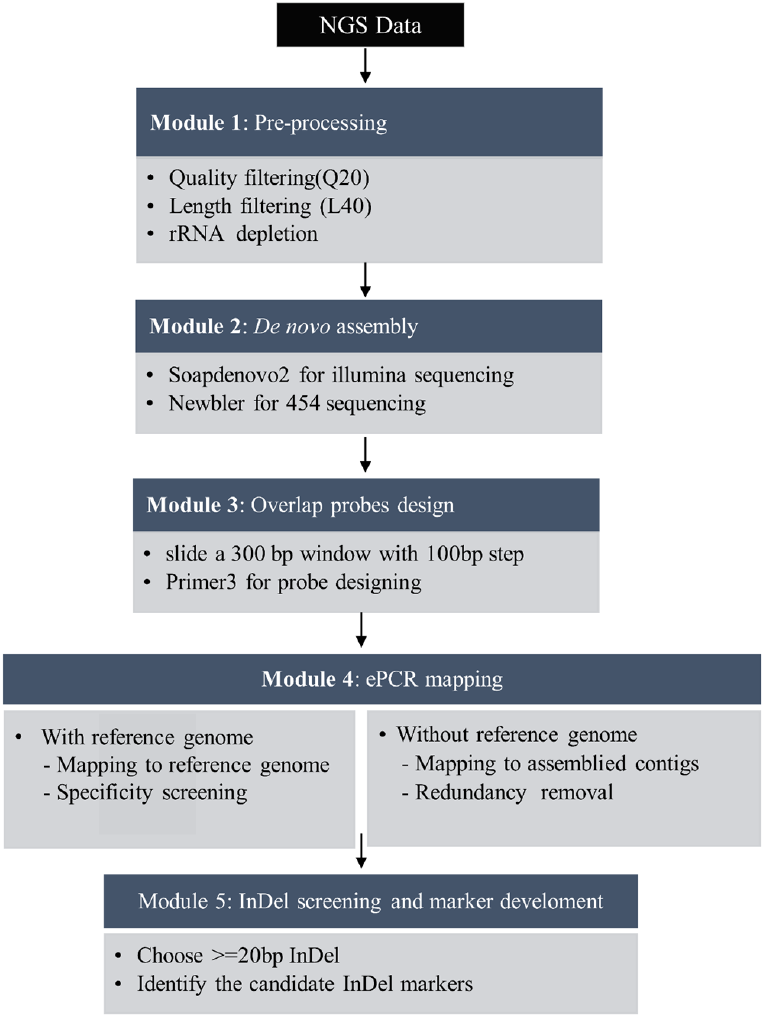
mInDel workflow. Graphic summary of mInDel workflow: the pipeline accepts NGS data as input and then proceeds automatically to perform several independent analyses, most of which can be selected or excluded according to the user’s needs. Module 1: Pre-process analysis. Module 2: De novo assembly (Illumina or 454 sequencing platform). Module 3: Overlap primer design. Module 4: ePCR mapping and specificity screening. Module 5: InDel screening and marker development.

### Pre-processing

This module is intended for quality control of sequence data, filtering against low quality, short reads and low complex regions. First, the Btrim [21] program is employed to remove low quality nucleotides at the 3’-end of a read by Q20 (Phred score of 20). Then, a custom Perl script removes sequence reads shorter than 40 bp (L40). When a sequence from a sequence-pair was removed, the remaining one is put into a separate file and used as a singleton during *De novo* assembly.

### *De novo* assembly

This module is designed for a high quality sequence assembly. We provide two different assembly strategies for two commonly used platforms (Illumina, Sanger and 454). For the Illumina platform with relatively shorter reads, we adopt the de-Bruijn graphs algorithm [22,23], which was specifically designed for processing short sequences and has been shown to be effective for sequence assembly. For Sanger or 454 platform with relatively long reads, the overlap algorithm [24,25] is employed for its efficacy at producing high quality assemblies of genome data. Here, SOAPdenovo2 [26] (Illumina Platform) or Newbler (sanger or 454 Platform) (http://www.454.com/products/analysis-software/) are used for sequence assembly using a coverage cutoff of 5 and discarding contigs shorter than 100 bp. By providing two different assembly strategies, mInDel enables users to choose flexibly according to their specificneeds.

### Overlap PCR Probe Design

This module is designed to generate high quality primer pairs. It generates large tracks of primer pairs, with estimated PCR products in overlap spanning the entire input region. This module firstly generates overlap fragments from the given sequence using a sliding window method. The overlap fragments are then passed on to the PCR primer selecting program Primer3 [27]. Primer3 then detects primer sets based on user’s setting criteria. The output file is parsed and generates a list of probe primers consisting of the primer sequence, the calculated melting temperature, the quality score of the primer set, primer positions, primer lengths, the PCR product length, and the amount of overlap between fragments.

### ePCR mapping and polymorphic screening

This module is designed for ePCR mapping. In silico PCR strategy is used to predict possible InDel differences from orthologous and homologous loci between any two genotypes. Primer sets generated from the previous module are run through in silico PCR analysis, with a threshold less than 3 mismatch bases using Bowtie aligner [28]. PCR products from a given region with different amplicon sizes are used to mine potential InDels among orthologous and homologous loci between the two samples. As a result, InDels larger than certain base pairs are preferentially mapped to the reference genome for single amplified locus screening. Finally, the site-specific InDel primer pairs are chosen.

### InDel marker development

In the final post-processing step, mInDel pipeline performs the post-processing to integrate core results and generate tab-delimited or Excel-compatible files. The final output compiles information for candidate InDel markers including forward and reverse primers, the product size (sample A and sample B), the InDel size (delta) and the chromosome location.

### Test data

A small set of test data is provided with mInDel. It consists of simulated Illumina 100bp *2 paired-end reads, and raw reference sequence. These data enable testing of most commonly used features. Users are strongly encouraged to run this test with the command ‘*sh test.sh*’ after installation which usually takes about 5 minutes to run on a 8-core 2.5 GHz computer. The test data also serves as the foundation for the example analysis described in the TUTORIAL file. We routinely run this test in the course of adding and refactoring code.

## Results and discussion

### InDel markers development between maize inbred lines

In addition to the small test data sets included with mInDel, we demonstrate here an example analysis of larger data sets from four maize inbred lines (B73 as the reference genome, Mo17, Qi319 and Zheng58). Mo17vsB73 and Qi319vsZheng58 were two experiments used for testing the mInDel pipeline performance under two different conditions: with and without the reference genome.

The Mo17, Qi319 and Zheng58 genome data was acquired from the NCBI’s Sequence Read Archive (SRA) database while the latest B73 reference genome data (V3) was from plant ensembl databases (http://plants.ensembl.org/Zea_mays). A total of 24.8 Gb 454 sequencing data from Mo17, as well as 30.4Gb and 12.5 Gb Illumina paired-end 100 bp reads for Qi319 and Zheng58 genotypes were retrieved from NCBI’s SRA database respectively.

The raw FASTQ-formatted reads were then pre-processed by Q20 and L40 filtering using mInDel’s quality control module. As a result, 21.2 Gb cleaned 454 reads from Mo17, along with 28.6 Gb and 10.1 Gb high quality Illumina reads from Qi319 and Zheng58 were obtained. The clean reads were then separately assigned into mInDel’s *de novo* assembly process. Contig assembly was performed with de bruijn graph method by SOAPdenovo for illumina sequences and with overlap method by Newbler for 454 sequences to ensure assembly quality. 1,351,472 contigs for Mo17, 476,470 contigs for Qi319 and 395,075 contigs for Zheng58 (>=100 bp) were generated and accounted for approximately 75%, 19.5% and 8.8% of the B73 reference genome. The contig N50 of the assemblies was 2,012 bp, 1,157 bp and 457 bp, with the longest contig being 33,817 bp, 18,068 bp and 7,286 bp for Mo17, Qi319 and Zheng58 respectively.

### Overlap PCR Primer Design

Furthermore, mInDel uses the open source Primer3 software to identify primers and probes with desired thermodynamic properties from candidate sequences in batch mode. At this stage of the pipeline, each contig larger than 300 bp was cut into several segments by sliding window of 300 bp with a step 150 bp, which ensured the overlapping amplified regions to cover the whole sequence. The amount of overlap is calculated from the 5’ end of a fragment to the 3’ end of the previous (adjacent) fragment and takes into account of the primer lengths. Generally, we have found that when sequencing these PCR products with dye-primer sequencing an overlap 150-200 bp is best. As a result, a total of 4,205,672 and 610,091 primer probes from Mo17 and Zheng58 were generated using mInDel.

### ePCR and polymorphic screening

In this study, two groups, Mo17vsB73 and Qi319vsZheng58, were assigned as two distinct data sets for all possible InDels loci based on in silico PCR analysis. Primer probes from Mo17 and Zheng58 contigs were run against the B73 reference genome sequences and Qi319 assembled contigs using in silico PCR analysis with a threshold less than 3 mismatch bases. In total, 1,855,222 primer probes from Mo17 were successfully mapped to B73 genome, while 307,345 from Zheng58 were mapped to Qi319 assembled contigs. Of mapped primers, 756,723 and 226,680 primer pairs were mapped to unique locations in B73 and Qi319 genomes and were considered as site-specific or single-copy primers, whereas primers mapped to multiple positions were discarded. Furthermore, a total of 739,250 and 223,084 InDel loci from group Mo17vsB73 and group Qi319vsZheng58 were identified using mInDel.

For InDel sizes, 133,918 and 44,389 markers have an InDel of 1-20 bp, while 594,717 and 178,695 markers have an InDel greater than or equal to 20 bp. The average length of the InDels was 71 bp and 66 bp, with the maximum InDels up to 387 bp and 371 bp. Finally, single-copy primer pairs with InDel size larger than 20 bp were selected and considered as candidate InDel markers for the ease of PCR detection and electrophoresis screening.

### Experimental validation of mInDel

To verify the accuracy and efficiency of mInDel, primer pairs randomly chosen across genome from each of the two comparison groups were synthesized for polymorphism detection between maize inbred lines Mo17 and B73 and between Qi319 and Zheng58. Of 318 sampling primers from Mo17vsB73, 276 (86.8%) showed polymorphisms with the expected amplicon sizes. In group Qi319vsZheng58, 437 of 602 InDel primer pair (72.6%) showed polymorphisms.

### Application on QTL fine mapping

Numbers of molecular markers are crucial to a high resolution genetic mapping. Marker saturation is one way to achieving fine mapping on recombination-rich regions with a yet saturated genetic map. Here, we successfully narrowed down one grain starch’s QTL region from a maize segregating population using InDel markers developed from mInDel pipeline. 90 InDel markers were developed for both regions by mInDel. As a result, 71 InDels showed polymorphisms between two parents, and 51 were successfully genetically mapped on regions with the synteny conservation to physical locations.

**Table 1:**
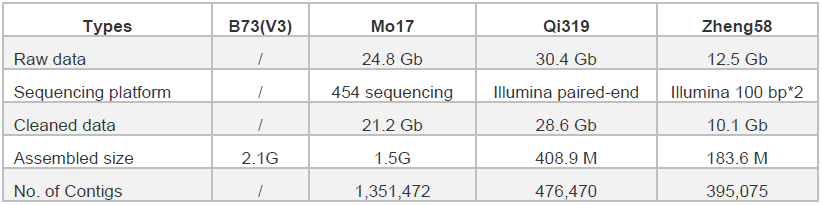
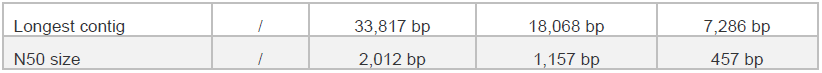
Summary of three genotypes genome data.

| Types | B73(V3) | Mo17 | Qi319 | Zheng58 |
| --- | --- | --- | --- | --- |
| Raw data | / | 24.8 Gb | 30.4 Gb | 12.5 Gb |
| Sequencing platform | / | 454 sequencing | Illumina paired-end | Illumina 100 bp*2 |
| Cleaned data | / | 21.2 Gb | 28.6 Gb | 10.1 Gb |
| Assembled size | 2.1G | 1.5G | 408.9 M | 183.6 M |
| No. of Contigs | / | 1,351,472 | 476,470 | 395,075 |
| Longest contig | / | 33,817 bp | 18,068 bp | 7,286 bp |
| N50 size | / | 2,012 bp | 1,157 bp | 457 bp |

**Table 2:** Summary of InDel markers developed from two groups.

| **Types** | **Mo17vsB73**<br>**(with reference genome)** | **Qi319vsZheng58**<br>**(without reference geome)** |
| --- | --- | --- |
| Total InDels | 739,250 | 223,084 |
| Mean size | 71 | 66 |
| 1~10bp | 76,376 | 25,144 |
| 11~20bp | 63,714 | 21,291 |
| 21~100bp | 374,847 | 123,788 |
| >100bp | 213,698 | 52,861 |
| sampling polymorphic rate | 86.8% (276 in 318) | 72.6% (437 in 602) |

## Conclusion

mInDel is an efficient and robust high-throughput pipeline in developing InDel markers between any two genomes. While most parameters work well with default settings, mInDel also offers great customization on parameter tuning. All modules are able to run independently, with the input/output features offering ample options and great flexibility. As a freely available, high-throughput and fully integrated software for InDel marker development, mInDel provides a useful resource for various genetics and genomics research and marker assisted breeding.

## Availability and requirements

**Project name:** mInDel

**Project home page:** www.github.com/lyd0527/mInDel

**Operating system(s):** Linux OS. Tests were performed in Centos and Ubuntu Linux systems. Some minor modifications are needed for other operation systems.

**Programming language:** Perl 5.10 or above, Shell.

**Other requirements:** Primer 3.2.0 or above and Bowtie any version.

## Abbreviation

InDel: insertion and deletion

NGS: Next Generation Sequencing

## Authors’ contributions

HZ and YDL conceived the study and wrote the manuscript. YDL developed the software for the analysis. XLZ and YHL compared the mInDel signatures with the experimentally verified signatures. All authors read and approved the final manuscript.

## Acknowledgements

This work was supported by the Natural Science Foundation of China(NSFC) under Grant No. 31271728, Jiangsu Agriculture Science and Technology Innovation Fund (JASTIF) under Grant No. CX(13)5055.

